# Hippocampal ripples linked to recognition memory judgments in children

**DOI:** 10.1101/2023.09.19.558300

**Authors:** Yitzhak Norman, Georg Dorfmüller, Delphine Taussig, Nathalie Dorison, Martine Fohlen, Mathilde Chipaux, Sarah Ferrand-Sorbets, Christine Bulteau, Michal Harel, Rafael Malach, Mikael Levy

## Abstract

Intracranial studies have demonstrated significant roles for hippocampal ripples in human declarative memory. Yet, the developmental trajectory and contribution of ripples to memory processes in children remain unknown. We studied hippocampal ripple activity using intracranial recordings in 14 children (age: 6-14) undergoing epilepsy monitoring. After watching a pink panther cartoon twice, participants engaged in an old/new recognition test, determining whether events portrayed in short (4s) videoclips stemmed from the cartoon they had just viewed. Our results reveal a significant rise in ripple rate during successful recognition of familiar events. An anticipatory decline in ripple rate preceded recognition errors. Interestingly, when participants viewed the cartoon passively during the initial full-length screenings, the overall temporal pattern of ripple activation remained consistent across repetitions, and significantly distinct from explicit recognition. We conclude that hippocampal ripples in children play a key role in declarative memory, supporting the explicit identification of previously encountered events.

## Introduction

The human brain undergoes significant structural and functional changes during childhood and adolescence ^1–4^. These changes occur both in large-scale cortical networks ^5,6^ and in more localized prefrontal and medial temporal lobe (MTL) structures that support memory functions ^7–13^. Neuroimaging studies on hippocampal development have shown a heterogeneous pattern of age-related changes across different hippocampal regions and subfields ^8,12,14–17^. Such structural and functional changes – which generally exhibit a complex, non-linear, developmental trajectory – play a role in mediating age-related changes in declarative memory capabilities^8,9,13,14,18,19.^

While these studies have provided valuable insights into hippocampal development and its impact on memory performance, there remains a gap in our understanding of how specific hippocampal mechanisms, that selectively contribute to declarative memory processing, mature with age. One such key mechanism, that received significant attention in recent years, is the extremely short and high frequency activity bursts termed hippocampal ripples. Ripples play a key role in various memory functions ^20,21^. Extant literature from animal studies has showcased their involvement in memory-guided decision making ^20,22–24^ and offline systems consolidation ^25–30^, while research in humans has associated them with retrieval of declarative memory ^31–37^.

However, despite the mounting evidence supporting the pivotal mnemonic role of ripples in the adult brain, it remains uncertain whether ripples exist in the hippocampus of young children and if they serve similar functions. Exploring ripple function across different age groups is crucial for comprehending the overall role of this vital hippocampal mechanism and how it evolves over time in relation to memory capabilities.

This study marks the first attempt to directly address these issues by investigating the role of hippocampal ripples in school-age children. Using a naturalistic cartoon-watching task and a non-verbal recognition test, we not only confirmed the presence and functional significance of ripples in the hippocampus of young children but also linked them with explicit familiarity judgments – an important yet unexplored aspect of ripple function in humans.

## Results

The study is based on intra-cranial recordings in 14 patients (8 females) ranging in age from 6 to 14 years (mean ± SD: 9.46 ± 2.6; see Extended Data Fig. 1 for additional demographic data). We focused on data obtained from 79 recording sites that were confirmed anatomically to be within the hippocampus (see Figure 1a-c and Methods).

**Figure 1.**
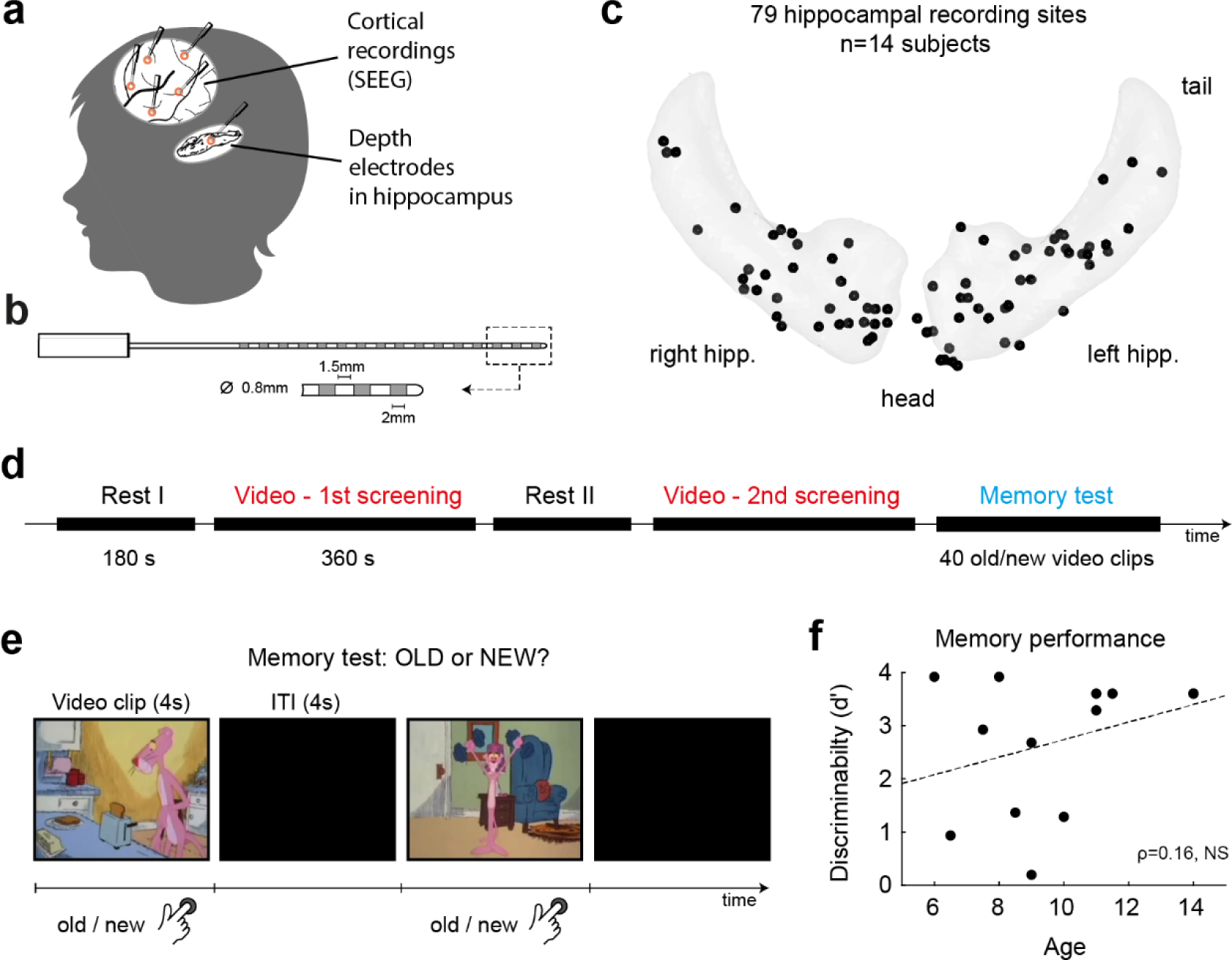
Experimental design, memory performance and intracranial recordings. (a) Each patient was implanted with multiple Stereo-EEG depth electrodes (SEEG) in the cortex and the hippocampus. (b) Schematic diagram of a typical SEEG depth electrode. (c) An overview of the electrodes’ anatomical location. Localization was done in the native space of each patient’s brain based on Freesurfer’s hippocampal subfield parcellation ^40^. For the ripples analysis, only electrodes located in CA1, CA2, CA3, or the subiculum were included, resulting in a total of 79 electrodes across 14 patients. (d) The task began with a 3-minute period of closed-eye rest, followed by a 6.5-minute viewing of a pink panther cartoon. This was followed by another rest period and a repeat viewing of the same cartoon. The recording session concluded with a recognition test. (e) During the memory test participants were presented with 40 distinct 4-second video clips. These clips either originated from the same pink panther episode or a different one. Using a mouse button, the participants indicated for each clip whether it was familiar (old) or unfamiliar (new). Screenshots from the Pink Panther cartoon are used under fair use for educational purposes. Copyright: THE PINK PANTHER™ & © 1964–2023 MGM. Memory performance (d’: *z*(*hit*) − *z*(*false alarm*)) among the participants was high (2.69 ± 0.36) and showed no correlation with age (Spearman’s ρ=0.16, NS). Each dot represents a single participant.

Briefly, the task began with a 3-minute resting period, after which the participants watched two consecutive screenings of a 6.5-minute audiovisual pink panther cartoon. The two screenings were separated by an additional 3-minute resting period (Figure 1d). Subsequently, the participants underwent an old/new recognition test, in which they were presented with short video clips lasting 4 seconds each. These video clips were extracted either from the original cartoon (n=20) or from a different pink panther episode revolving around a similar theme (n=20; see Figure 1e and Methods for further details). The participants were asked to indicate for each video clip whether it was old or new to them via a button press.

The participants performed the task successfully, achieving an overall accuracy of 86.4 ± 4.9%. Memory performance did not correlate with age. Age did not significantly predict accuracy (Spearman’s ρ=0.25, p>0.41) nor discriminability (d-prime) (ρ=0.16, p>0.59; Figure 1f), which aligns with previous reports on recognition performance in similar age groups ^38,39^.

### Characteristics of hippocampal ripples in children

Throughout the recording session, iEEG signals were acquired from the hippocampal electrodes, and ripple events were detected following standard protocols ^41^ that closely resemble the methods described in our earlier studies ^35,42^ (see Methods). Examples of ripple detection are depicted in Figure 2. Notably, the detected events showed a remarkably stereotypical and consistent spectral signature characterized by a brief temporal duration and a prominent increase in oscillatory power concentrated at ∼90 Hz, which reflects the intense, coordinated, multi-neuron activity of hippocampal ripple events (Figure 2c-e) ^43,44^.

**Figure 2.**
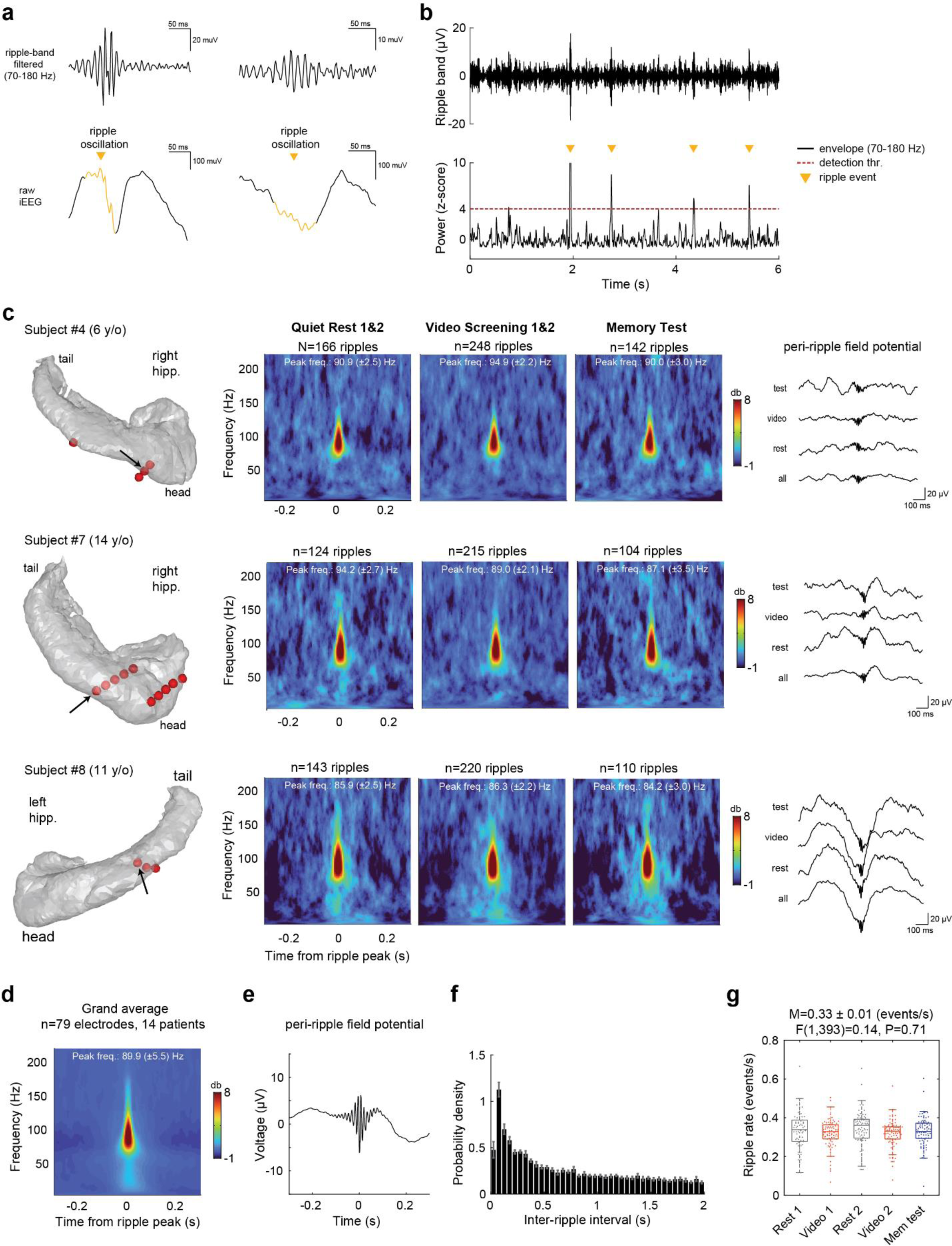
**Hippocampal ripples detection and their characteristics in children.** (a) Two traces of representative ripple events detected in one patient. (b) Ripple detection procedure, showing examples of four candidate ripples (marked by orange triangle) as they appear in a CA1 recording site. (c) Peri-ripple spectrograms and average field-potentials in representative CA1 recording sites in three patients (6, 14 and 11 years old). The left panel shows the anatomical location of each electrode within the hippocampus. Note the similar spectral signature of ripples present in these three patients, despite their varying ages. Ripples’ peak frequency, duration and amplitude remain stable across the different experimental conditions: quiet rest, watching a full-length cartoon, and the memory test. (d-e) Grand-average peri-ripple spectrogram and field-potential, computed across all hippocampal electrodes, in 14 subjects. (f) Grand-average inter-ripple-interval distribution, in 50 ms bins. The basal (mean) ripple rate remained stable throughout the experiment, with no significant differences between task phases. However, a more transient change in ripple rate did occur during the memory test, as will be demonstrated later.

Specifically, the hippocampal ripples recorded exhibited a peak frequency of 89.9 ± 5.5 Hz (Figure 2d) and had an average duration of 34 ± 6.4 ms. Ripple duration was determined as the time interval between the onset and the offset of the ripple, that is, when the power of the oscillatory event fell below a 2σ threshold (see Methods). The overall ripple rate throughout the experimental duration was 0.33 ± 0.01 events/sec, demonstrating a typical inter-ripple interval distribution (Figure 2f-g). For comparison with typical ripple properties in adults see ^35,36,42^.

Applying a linear mixed-effects model to the individual electrode data, showed no significant differences in basal ripple rate, ripple duration, or ripple amplitude across the various experimental conditions (rate: F(1,392) = 0.14, P = 0.71; duration: F(1,392) = 3.59, P = 0.06; amplitude: F(1,392) = 0.06, P = 0.8), nor significant effects of age. This further emphasizes the stability of the observed stereotypical ripple properties across the participants and throughout the experiment ^45^. The formulation of the mixed-effects models employed was as follows:

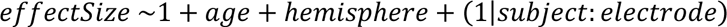

where x is the measure of interest in each analysis (i.e., ripple rate, duration, or amplitude).

### Ripple activity linked to recognition memory

Examining ripple rate modulations during the two movie repeats and the old/new recognition test revealed several significant findings that link ripple activation with recognition memory (Figure 3).

**Figure 3.**
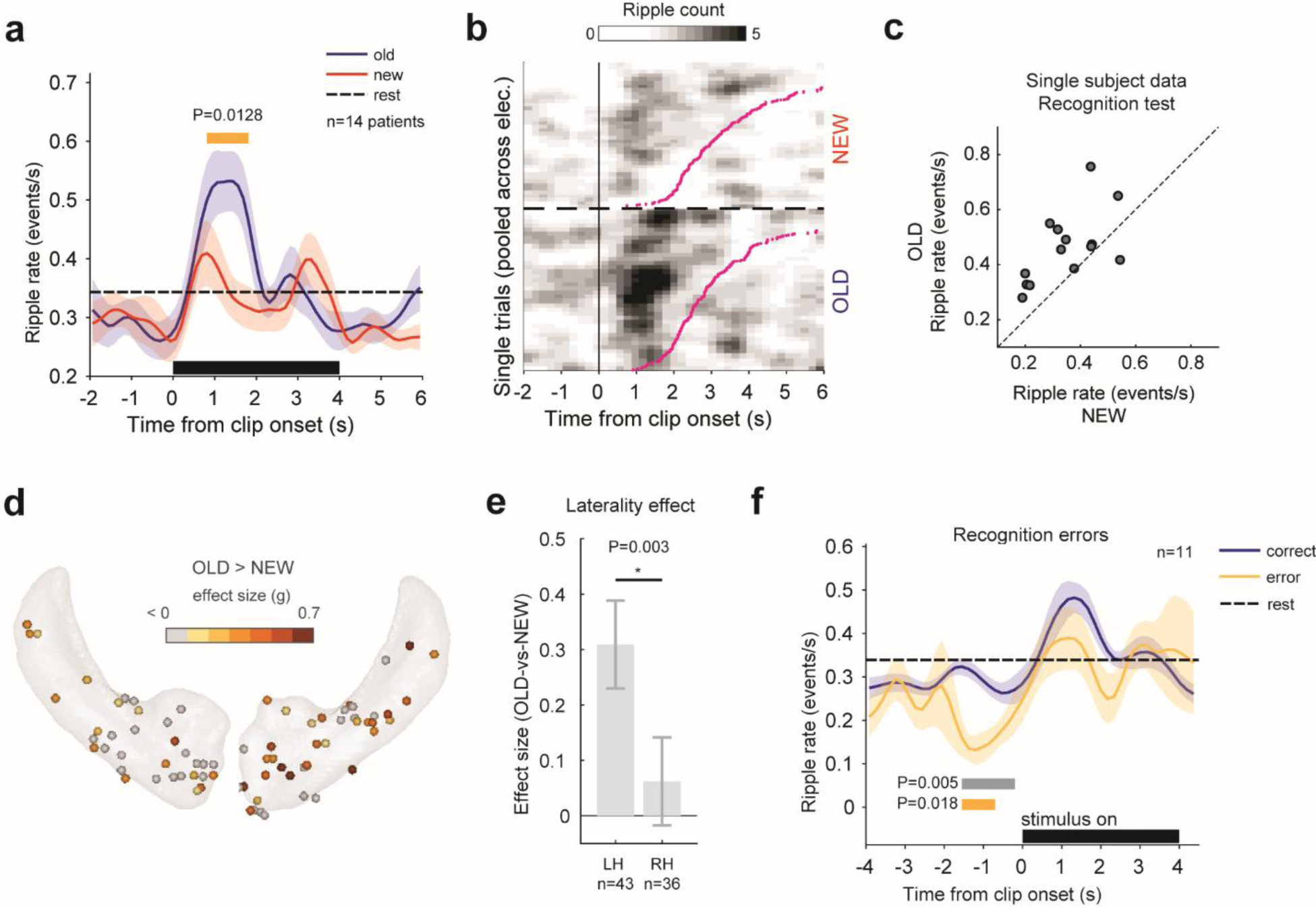
**Ripples activation linked to recognition judgment** (a) Peri-event time histogram (PETH) aligned to the videoclip onset, showing an instantaneous ripple rate increase during the recognition test. Participants viewed 4-second videoclips extracted either from the pink panther cartoon they watched (“old”, n=20 clips) or from another pink panther cartoon (“new”, n=20 clip), in pseudorandom order. When subjects recognized a clip as familiar, there was a momentary increase in ripple rate (P=0.0128, n=14 patients, cluster-based permutation test). (b) Hippocampal ripple density plot showing the increase in ripple probability following the presentation of a familiar (“old”) clip. Trials were stacked together across all electrodes and sorted according to reaction time (pink curve). Ripples’ density was smoothed using a 5-bins-wide 2d Gaussian filter for visualization purposes. (c) Scatter plot of ripple rate in individual patients, showing “old” versus “new” clips. Except for one patient (right- hemisphere implant), all patients demonstrated a higher ripple rate when recognizing a familiar clip. (d) Anatomical distribution of electrodes showing an old>new effect. Electrodes in the left hippocampus overall showed a stronger effect. (e) A mixed-effects model applied to the data revealed a significant hemisphere effect, indicating more robust recognition-related ripple activation in the left hippocampus compared to the right. (f) PETHs of correct versus incorrect trails, showing a momentary reduction in ripple rate anticipating recognition errors (errors vs. rest, P=0.005 (grey bar); errors vs correct trials, P=0.018 (orange bar); cluster-based permutation test).

First, we discovered a stimulus-driven, transient, increase in ripple rate. This effect was found by computing a peri-event time histogram (PETH) of ripples, aligned to the onset of videoclips in the recognition test. We determined the optimal bin size using Scott’s method^46^ and smoothed the PETH using a 7-point triangular window.

The PETHs indicated a transient increase in ripple rate a few hundred milliseconds (half-height: 588 ± 53 ms; peak: 1015 ± 33 ms) after the onset of the videoclip (clip-vs-rest: P=0.016, N=14 patients, cluster-based permutation test). This increase was evident for both clips judged as old or new (Figure 3a) and could also be observed at the single trial level, as depicted in Figure 3b.

Second, a significant enhancement in ripple rate was observed when the presented video clip was *explicitly* judged as familiar (recognized as ‘old’) compared to unfamiliar (judged as ‘new’). This enhancement is clearly demonstrated by comparing the blue and red curves in Figure 3a (P=0.0128, n=14 subjects, cluster-based permutation test), and by contrasting the ripple rate elicited during old and new clips individually for each subject (Figure 3c). For the latter, ripple rate was averaged over a time window of 2 s centered on the peak of the grand average response reported above (computed across all clips, old and new).

Third, the observed effect exhibited a hemispheric difference, with a significantly more prominent manifestation in the left hippocampus compared to the right hippocampus. This difference is illustrated visually at the single electrode level in the old-vs-new effect size map shown in Figure 3d and statistically at the group-level in Figure 3e. In this analysis, the ripple rate was individually computed for each site using a fixed time window of 2s centered on the peak of the grand average ripple rate response. For the statistical comparison, we fitted a linear mixed effects model to the old-vs-new effect size values (Hedges’ g), formulated as follows:

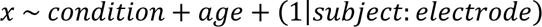

This analysis revealed a significant hemisphere effect (t(76)=3.09, P=0.003). Importantly, it also indicated the absent of an age effect (t(76) = 0.002, P = 0.96), suggesting that the recognition signal mediated by ripples is already fairly established at this school-age range. Notably, this finding aligns well with the estimated rate of neuronal maturation in the tri-synaptic circuit in humans, which is believed to reach completion before the age of 5 ^47,48^.

An interesting question concerned the link of ripple rate to errors made. Examining ripple rates during trials where participants made recognition errors – i.e., failed to recognize an old clip or incorrectly identified a new one as old (13.6 ± 4.9% of trials on average) – we found a brief decrease in ripple rate occurring 1-2 seconds prior to the recognition error. This reduction was statistically significant when compared to both correct trials and resting state conditions (errors vs correct trials, P=0.018; errors vs. rest, P=0.005; N=11 patients, cluster-based permutation test; see Figure 3f).

Interestingly, in stark contrast to the significant increase in ripple activation during the explicit recognition judgment, the observed ripple rate responses for the same animated events when viewed passively during the original full-length screenings (i.e., outside of the context of a recognition test), showed no enhancement (Figure 4a), neither during the initial nor the repeated screenings. This finding suggests that the rise in ripple rate during the recognition test was specifically associated with the explicit, intentional, memory judgment being made, rather than the passive perception of the repeated dynamic visual content itself.

**Figure 4.**
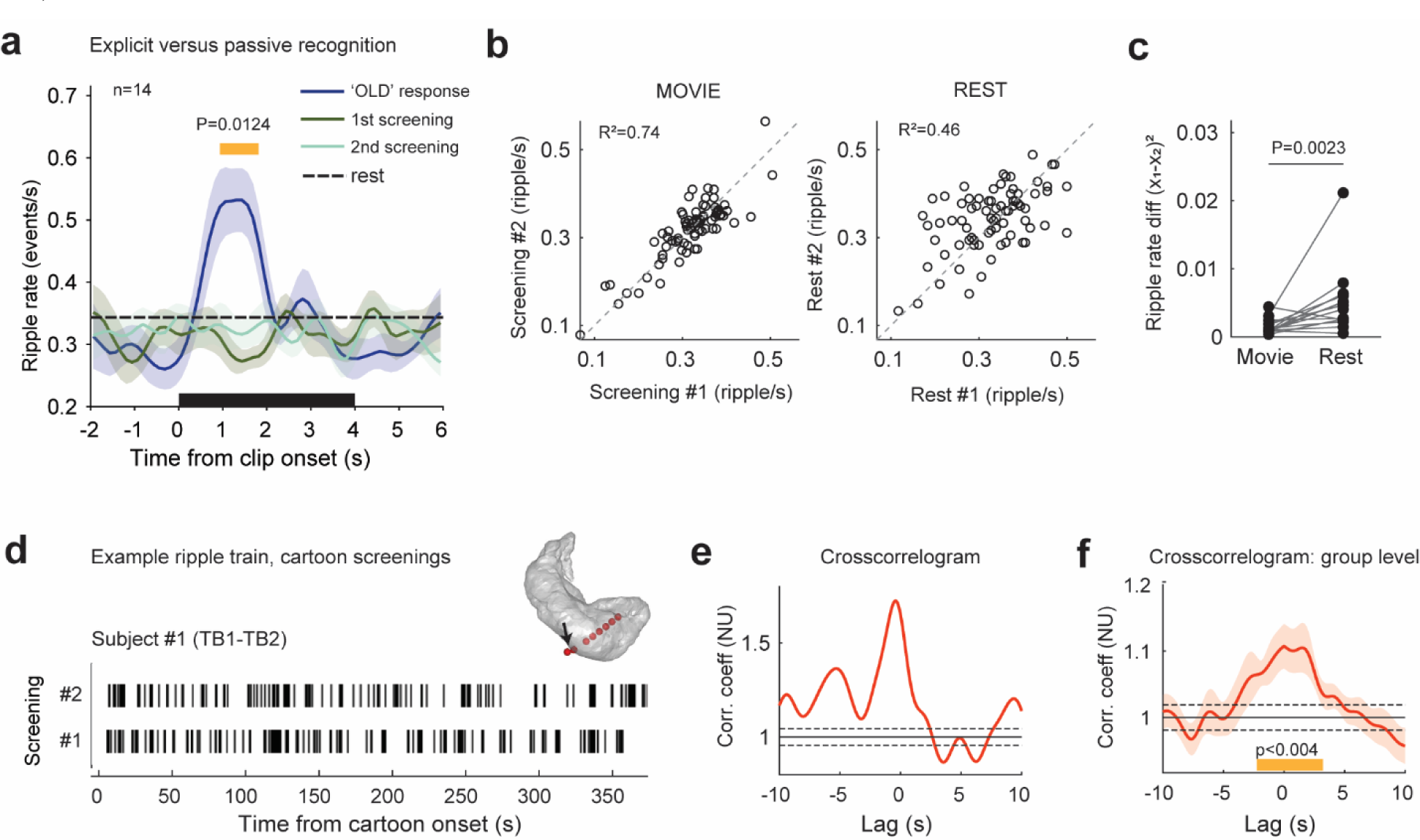
**Ripple activation during explicit versus implicit recognition** (a) Ripples PSTHs showing that when the same animated events were viewed passively during the full-length screenings, neither the initial screening (green) nor the repeated screening (turquoise) exhibited an increase in ripple rate. Thus, the increase in ripple rate during the recognition test was specific to explicit memory judgments. (b) Left: Ripple rate correlation between the first and second full-length screenings (R2=0.86). Right: Ripple rate correlation between the first and second resting periods (R2=0.46). Note the striking stability of ripple rates across the two screenings of the cartoon. (c) Directly comparing the stability of ripple rates during the repeated cartoon screenings against that observed during the repeated resting periods revealed significantly higher stability during the cartoon repetitions (P=0.0023, n=14 patients, signed-rank test). Stability was quantified by the squared difference between the two repeats, where lower values indicate higher stability. The differences were average across electrodes, individually in each patient. (d-e) An illustrative example of ripple train during the two screenings of the cartoon in one recording site (see arrow for electrode location) and their cross-correlation. Note the prominent peak around zero lag, highlighting the consistency in the temporal activation profile of ripples during the cartoon. (f) Mean cross correlation between the first and second screenings of the cartoon, indicating a significant peak around zero lag (P<0.004, N=14 patients, cluster-based permutation test). The black dashed line represents the mean±SD of cross-correlation magnitudes when ripple trains were randomly shifted in a circular manner 50 times.

While the observed momentary rise in ripple rate pertained specifically to moments of explicit recognition judgment, it was crucial to investigate how the repetition of the full-length cartoon and its accompanying cognitive processes influenced the overall pattern of ripple activation during passive viewing. Notably, as shown in Figure 2g, re-watching the same cartoon did not yield an overall increase in ripple rate. However, when analyzing the correlation between ripple rate elicited during the first and second screenings across recording sites, a distinct observation emerged: encountering the same cartoon for a second time involved the recurrence of a similar pattern of ripple activation across electrodes (Figure 4b). This overall pattern stability was then compared to that observed during the two resting state periods (measured immediately before each cartoon screening), confirming that the overall activity pattern during the cartoon was significantly more stable compared to the one emerging during the resting state condition (Figure 4c; P=0.0023, N=14 patients, signed-rank test comparing the sum of squares of ripple rate differences between the two screenings with that of the resting state periods).

To further explore whether the above stability was also reflected in the temporal pattern of ripple activation, i.e., the timing of the ripples, we conducted a cross-correlation analysis between the ripple train from the initial and repeated screenings at each recording site (see Methods for more detail; an example crosscorrelogram is depicted in Figure 4d-e).

The analysis unveiled a noticeable peak around zero lag, indicating that ripple activations occurred consistently around similar times during the movie (Figure 4f; P<0.004, N=14 patients, cluster-based permutation test). Hence, the re-experience of the same cartoon resulted in a similar pattern of ripple activation over time. It is worth noting that the cross-correlation peak showed a rather broad and gradual pattern, suggesting that ripple activation might not specifically represent precisely timed sensory events. Instead, it could indicate recuring cognitive or mnemonic processes that are associated with the viewing experience but less tightly bound to a precise moment in time.

To assess whether these temporal activation profiles reemerged also during the recognition test, we isolated intervals from the full-length cartoon that aligned with the videoclips used in the memory test. Within these intervals, we recalculated the cross-correlation between the first and second screenings and between each screening and the memory test. While the results replicated the cross-correlation found between the two initial screenings, we did not observe a significant correlation between the ripples elicited during the explicit recognition and those elicited during the passive viewings (Extended Data Fig. 2). This suggests that the momentary rise in ripple activity during the explicit recognition judgment constitutes a distinct hippocampal process, characterized by both an elevated ripple rate and a different activation profile. This process likely contributes to an explicit sense of familiarity, aiding in the conscious recognition of previously encountered events.

### Ripples rate differs from HFB signal

Finally, to further investigate the underlying processes, we examined whether the recognition effect was specific to ripple activity or also evident in other measures of local neuronal activity, particularly the high-frequency broadband power (HFB: 60-128 Hz; also known as high-gamma; see Figure 5).

**Figure 5.**
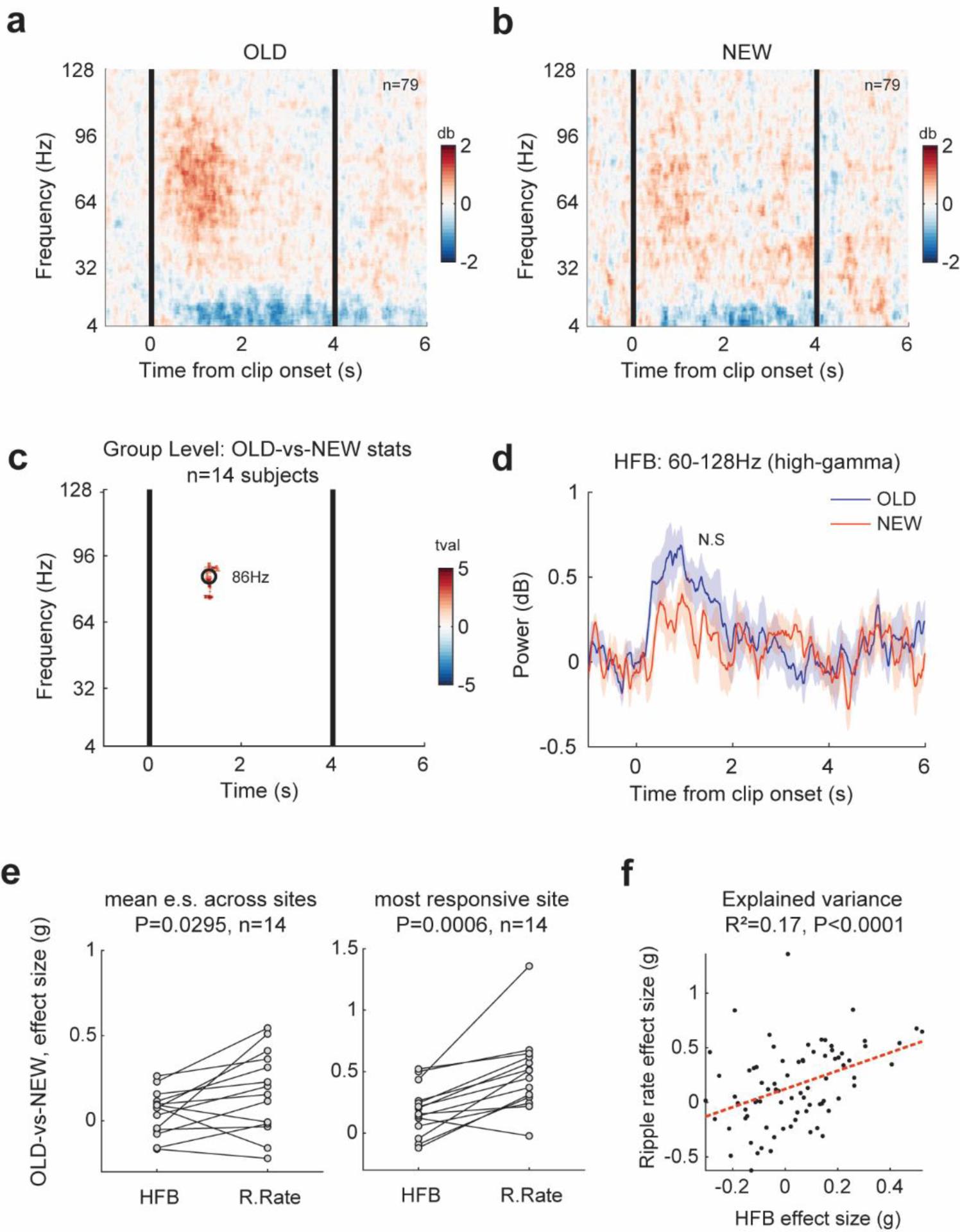
**Ripple rate shows greater sensitivity to the mnemonic process compared to the High Frequency Broadband (HFB) signal.** (a-b) Grand average spectrogram compute across all hippocampal sites, showing high-frequency broadband activity during clips correctly recognized as OLD (a) and clips correctly identified as NEW (b). (c) Statistical comparison between the OLD and NEW spectrograms indicated a single significant cluster concentrated within the ripple frequency band, specifically between 82-94 Hz (N=14, cluster-based permutation test; semi-transparent time-frequency bins indicate non-significant clusters). (d) To directly examine the mnemonic specificity of the high-frequency broadband (HFB) response, we averaged the spectrogram rows within the high-gamma band (60-128 Hz) and compared the responses between correctly recognized OLD and NEW clips. Our analysis revealed no statistically significant differences between the two responses (P=NS, n=14 patients, cluster-based permutation test). (e) Left: A direct comparison of effect sizes between the high-frequency broadband (HFB) signal and ripple rate for the ‘OLD-versus-NEW’ contrast, averaging across all hippocampal sites in each patient, demonstrated a significantly greater effect size for ripple rate compared to HFB (P=0.0295, n=14, Wilcoxon signed-rank test). Right: comparing the effect sizes obtained from the best HFB site and the best ripple-rate site in each patient, again showed a significantly larger effect size for ripple rate (P=0.0006, n=14, Wilcoxon signed-rank test). Correlation between the effect sizes obtained from HFB and ripple rate signals.

For the analysis we computed clip-triggered spectrograms (see Methods for detail) individually for correctly recognized old and new clips. Consistent with the ripple rate response described above, there was a general increase in high-frequency power following clip onset. A cluster-based permutation test indicated a sole significant cluster emerging ∼1.1 s following clip onset and concentrated within the ripple band, specifically between 82-94 Hz (P<0.05, N=14, cluster-based permutation test with 2000 iterations^49^). The timing of this cluster, which seems consistent with a previous iEEG study in adults^50^, aligns closely with the peak of the ripple rate response shown in Figure 3a. This implies that the power increase within this frequency range is likely a reflection of ripple activity.

Yet, for a more targeted examination of the high-frequency broadband (HFB) signal, we averaged the spectrogram rows within the high-gamma band (60-128 Hz) and compared the correctly recognized old clips with new ones. As expected, the HFB response displayed a general increase during clip presentation. However, unlike the ripple rate, the HFB response demonstrated less sensitivity to the memory process, showing only a minor increase in activation for old clips compared to new ones, which did not reach statistical significance (P=NS, n=14 patients, cluster-based permutation test; compare Figure 5d with Figure 3a).

Likewise, a direct comparison between the old-vs-new effect size of the ripple rate and the HFB power within the 4s duration of the video clips revealed a significantly stronger recognition effect for the ripple rate (P=0.0295, n=14, Wilcoxon signed-rank test). Even when comparing the single most recognition- selective HFB site in each patient to the single most recognition-selective ripple rate site in each patient (Figure 4e), the effect size was significantly larger for the ripple rate (P=0.0006, n=14, Wilcoxon signed-rank test). This analysis suggests that hippocampal ripples provide a more accurate reflection of the underlying mnemonic process compared to the overall population firing rate represented by HFB activity.

Lastly, considering that HFB and ripple rate are neural measurements involving semi-overlapping frequency bands, it became important to quantify not only the difference between these two signals but also to assess their association. To accomplish this, we computed the correlation between the effect sizes of ’old-vs-new’ contrasts in HFB and ripple rate across all hippocampal electrodes. This yielded a statistically significant correlation (R=0.42, P<0.0001, n=79 electrodes; Figure 4f). However, the magnitude of this correlation implies that broader alterations in hippocampal HFB power could explain no more than 17% of the variability in ripple rate during the recognition process, further emphasizing the functional distinction between these two signals.

## Discussion

The present results demonstrate a significant role for hippocampal ripples in recognition memory. Our findings both confirm and extend our previous reports of the involvement of hippocampal ripples in human memory function along two major lines. First, they extend these findings to the domain of brain development during childhood and early adolescence (ages 6-14). Second, they extend our analysis to the domain of naturalistic stimuli – watching a movie – shedding light on the distinct functions of ripples in memory processes encountered in daily experiences.

Our results highlight the cognitive specificity of ripple activation observed in young participants when viewing animated events passively in a cartoon, as opposed to viewing them as part of an explicit recognition test. The selective increase in ripple rate during the recognition test suggests that ripple activity is not merely a passive sensory response to the momentary visual content of the movie. Instead, it seems closely tied to the contextual situation and cognitive task in which this content is embedded.

Two potential factors, not necessarily mutually exclusive, may account for this difference in ripple activation. The first possibility could be the switch from a continuous, ongoing processing during the movie presentation to a discrete event-like presentation of an isolated clip. Indeed, this possibility is compatible with previous research emphasizing the importance of abrupt cuts and event boundaries in hippocampal function^51,52^. The second potential factor could be attributed to the cognitive task shift – from passive re-experiencing of movie content to an explicit recognition task that requires active engagement in declarative memory judgments to discern whether the presented movie event was familiar or new.

Importantly, the observation that recognition errors were associated with a notable reduction in ripple rate prior to the clip onset (Figure 3f) significantly supports the second explanation. It disentangles the association between ripple rate and the abrupt visual presentation while preserving a significant connection to the subsequent memory judgment. Therefore, considering these two observations together, it suggests that the rise in ripple rate observed during the memory test was specifically tied to the memory judgment.

Another, related, finding is the significant enhancement in ripple rate when patients were exposed to familiar movie clips compared to novel ones when performing the recognition task. This enhancement suggests that beyond the discreteness of the event and the overall decision-making task – there was a dimension of activation specifically related to the memory of the clips. Thus, this selectivity of ripple activation suggests that the hippocampal ripples are specifically involved in the active recall of previously stored materials as opposed to recognition attempts or mere processing of novel material. This interpretation is nicely compatible with earlier studies demonstrating ripple activation when adult participants perform autobiographical memory judgments ^37^, free recall of pictures ^35^ and words ^53^, and retrieval of paired associations ^54–56^. Importantly, these studies have shown that ripple activations were specifically linked to the retrieval process and were not observed when other cognitively effortful tasks were employed (e.g., arithmetic calculations^37^).

The substantial coverage and number of hippocampal recording sites (79 electrodes) allowed a coarse analysis of anatomical localization of the recognition effect. Our results did not show evidence for local clustering of preferential activation to old vs. new content within the hippocampus (Figure 3D). Instead, the results emphasize the distributed nature of hippocampal mnemonic representation, echoing a similar observation previously noted in autobiographical and semantic memory recall studies ^42^. This finding also aligns well with single-neuron studies in humans describing a fairly balanced mixture of hippocampal neurons preferentially activated by novel stimuli and those activated by repeated stimuli ^57^.

Yet, when comparing the hemispheric preferences – i.e., left versus right hippocampus – a significant left-hemisphere dominance was observed, aligning with our previous observations ^42^, as well as with a previous fMRI study showing a left-lateralized subsequent memory effect in children of a similar age tested on words^39^. Although the memory test we employed was non-verbal in nature, it is still interesting to consider whether the observed left-hippocampus preference might be connected to the language dominance of the left hemisphere. However, further exploration of this notion would necessitate a larger dataset and the inclusion of both verbal and non-verbal conditions ^58^.

During the initial screenings of the full-length cartoon, we observed a significant increase in the stability of ripple activation patterns compared to the resting state (Figure 4b). This stability manifested in both the overall spatial pattern of ripple activity – recording sites across the two hippocampi maintained a remarkably similar activity level during the first and second screenings – and in the timing of ripple activation. The cross-correlation between the ripple trains from the two screenings revealed a significant peak approximately 5-10 seconds in width, indicating that the reinstated temporal pattern across the two screenings was relatively slow, suggestive of gradual cognitive processes associated with the animated content. If the ripples were more precisely linked to the sensory content itself, one would expect a sharper cross-correlation function.

High-frequency broadband (HFB) power has been established as a reliable index of local neuronal population activity in cortical systems^59–63^. However, far less is known about this collective neuronal activity marker in hippocampal neurophysiology. It should be noted that the HFB power modulation in the cortex is likely to stem from the functional clustering of neuronal response properties in the well-established cortical columns. The fact that no such clustering has been documented in the hippocampus – poses a challenge in interpreting the precise neuronal basis of HFB power in this region. Furthermore, it remains unclear how much information overlap exists between hippocampal HFB and ripple rate – given their derivation from similar, partially overlapping frequency bands in the local field potential (ripples: ∼80-140 Hz, HFB: ∼60-160Hz).

Indeed, our analysis revealed some overlap between the memory-related ripple rate changes and concurrent HFB power modulation (see Figure 5f). However, importantly, the ripple rate changes showed a significantly higher level of mnemonic specificity – i.e., when comparing the preferential activation to old vs new content (Figure 5e). A likely explanation for these results is that the HFB and ripple rate may share a wide-spread, arousal-like, physiological modulations – leading to highly distributed increase in firing rates. However, specific contents such as the difference between old and new materials – may preferentially generate ripples while remaining unselective in terms of global neuronal activity in the hippocampus.

## Conclusions

Our findings provide the first evidence for the involvement of hippocampal ripples in declarative memory processes in school-age children. The results indicate that the electrophysiological characteristics of hippocampal ripples ^35,42^ appears early in the developing brain. This was demonstrated in the ripple dynamics, ripple frequency, functional selectivity, and mnemonic specializations. Specifically, our findings highlight the early development of cognitive specificity in ripple activation, which shows notable selectivity towards explicit memory judgments in young children. This reinforces the notion that hippocampal ripples during the waking state are closely tied to mnemonic processes that are consciously accessible and reportable ^32,35,36,42,54^, as opposed to the more automatic, sensory-driven components, that contribute to the sense of familiarity in a more implicit manner ^64^.

## Acknowledgments

We express our gratitude to the patients and their families for their generous willingness to participate in this study. This work was supported in part by the Zuckerman STEM Leadership Program (YN) and by the CIFAR Tanenbaum Fellowship (RM).

## Author contributions

YN, RM and ML conceived the study. YN designed the experiment. YN analyzed the data. RM and ML supervised the analysis. ML, SFS and GD performed the surgeries. YN and ML installed the experimental setup. ML ran the experiment and supervised all aspects of data collection. MF and CB performed neurological and neuropsychological evaluations. DT, ND and MC monitored and analyzed epileptiform activity. YN, MH, and ML contributed to data curation and electrode localization. RM secured funding. YN and RM wrote the paper. ML further contributed to the writing by reviewing and editing the manuscript.

## METHODS

### KEY RESOURCES TABLE

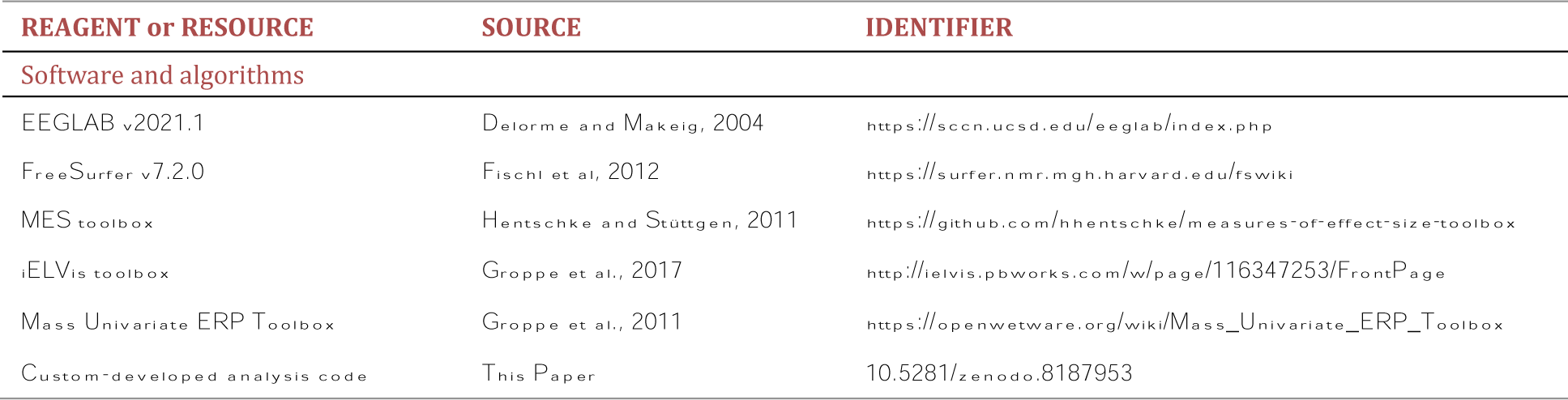

### Resource availability Lead contact

Further information and requests for resources should be directed to.

### Materials availability

This study did not generate any new reagents.

### Data and code availability

The data and code that support the conclusions are available upon reasonable request from YN and will be made publicly available on zenodo upon publication (DOI: 10.5281/zenodo.8187953).

### Experimental Model and Subject Details

#### Participants

14 children (8 females; age: M=9.4, range: 6-14; see Extended Data Fig. 1 for additional demographic details) with medicine-resistant epilepsy were implanted (G.D, S.F and M.L) with intracranial stereo-electroencephalography (SEEG) electrodes (DIXI® and ALCIS®) as part of their pre-surgical evaluation at Rothschild Hospital, Paris. Electrode implantation was performed using ROSA (Zimmer©) under general anesthesia, regular anti-epileptic drugs, and antibiotics regimen. The location of the implanted electrodes was done based on the pre-surgical hypothesis and confirmed post-operatively by computed tomography (CT). The patients included in this study were selected based on electrode coverage in the hippocampus. Each patient was monitored in the hospital for 5 days on average (4-15 days). All patients had left hemispheric dominance for language, as determined pre-surgically using functional MRI and/or SEEG language mapping. The study was approved by the Institutional Review Board (TNP-SEEG protocol V1-0 20151012 1/6), and a written informed consent to participate in the study was obtained from the child’s parents before surgery.

## METHOD DETAILS

### Intracranial EEG recordings

All electrophysiological data were recorded using a Micromed (Treviso, Italy) clinical monitoring system. Sampling rate was commonly 512 Hz (256 Hz in five patients) with a bandpass filter of 0.7-300Hz. None of the patients included in the analysis experienced any seizures throughout the experimental duration.

### Experimental Task

The task began with a 3-min period of closed-eyes rest, after which the participants watched a 6.5-minute audiovisual pink panther cartoon titled “Pink Breakfast”. After an additional 3-min resting state, participants re-watched the same cartoon. Upon completion of the second screening, the participants underwent an old/new recognition test. In this test, 40 short video clips, each lasting 4 seconds, were presented with no sounds. The clips were extracted either from the original cartoon (n=20; “Pink Breakfast”) or from a different episode with similar food themes (n=20; “Dietetic Pink”). The participants were instructed to indicate whether each clip was old or new by pressing the left/right mouse buttons as soon as they had an answer. Participants that were unlikely to execute the button press in a timely manner were instructed to respond verbally instead, and the experimenter pressed the button on their behalf (a total of 6 patients). Throughout the task, the parents of the participants remained quietly at their bedside.

### Electrode localization

Structural MRI were acquired using a (GE Healthcare 3 tesla SIGNA scanner) at the Rothschild Hospital. A preoperative T1-weighted 1mm isometric scan was co-registered to the post- operative CT through the procedure described previously ^42^, which is also extensively documented in the iELVis toolbox ^65^. Individual recoding sites were identified visually on the co-registered CT, and were marked in each subject’s preoperative MRI native space, using BioImage Suite ^66^. After processing the MRI in Freesurfer ^67^ to segment the cortex and hippocampus ^40^, we superimposed the hippocampal electrodes onto a 3D reconstruction of the hippocampal formation, allowing us to determine the precise subfield localization of each recording site within the native space of the patient’s brain.

### Preprocessing and data analysis

Data analysis was performed in Matlab 2021a (MathWorks Inc., Natick, MA) using EEGLAB v2021.1 ^68^ and custom-developed analysis code. The raw iEEG signal was examined statistically to identify and exclude noisy or corrupted channels from further analysis. Channels with voltage values, voltage derivatives, or RMS in the 99th percentile that were 5 standard deviations greater than the rest of the electrodes were marked for exclusion after visual inspection in the time and frequency domains. Preprocessing included converting all healthy iEEG channels into bipolar derivations. Electrodes in the hippocampus were referenced relative to a nearby white-matter contact, that was identified anatomically using FreeSurfer’s segmentation ^67^. The obtained bipolar signals were then notch-filtered to remove 50 Hz power line interference using a zero-lag linear-phase Hamming-windowed FIR band-stop filter (3 Hz wide).

### Hippocampal Ripples Detection

To detect hippocampal ripples, a slightly modified version of the method described in ^42^ was employed. This detection was conducted on electrodes located within <2 mm from the hippocampal subfields CA1, CA2, CA3 or subiculum, i.e., the main stations on the hippocampal output pathways where physiological ripple oscillations are most prominently observed ^69,70^. We delineated and reconstructed the individual hippocampal subfields based on the structural MRI scans of the patients ^40^. Prior to ripple detection, a reference signal from a nearby white-matter contact was subtracted to eliminate common noise. The iEEG time series was then filtered between 70-180Hz (using a zero-lag linear-phase Hamming windowed FIR filter), and the instantaneous analytic amplitude was computed using a Hilbert transform. For the 5 patients with a sampling rate of 256 Hz, a narrower ripple-band filter between 70- 128Hz was utilized. Based on our previously published procedure ^42^, extreme values were robustly estimated using Least-Median-Squares (LMS) ^71^ and clipped to 4 SD above the mean to minimize ripple-rate induced biasing. The clipped signal was then squared and smoothed (Kaiser-window FIR low-pass filter with 40 Hz cutoff), and the mean and SD were computed across the entire experimental duration to define the threshold for event detection. Events from the original (squared but unclipped) signal that exceeded 4 SD above baseline were selected as candidate ripple events. Event duration was expanded until ripple power fell below 2 SD. Events shorter than 20ms or longer than 200ms were excluded. Adjacent events with less than 30ms separation (peak-to-peak) were merged. Finally, ripple peaks were aligned to the trough (of the non-rectified signal) closest to the peak power.

A control detection was performed on the common average signal computed across all iEEG channels. Hippocampal ripple events that coincided with common average ripple-band peaks were removed, thus avoiding erroneous detection of transient electrical and muscular artifacts that tend to appear simultaneously on multiple channels ^41,72^.

To distinguish between physiological ripples and other high-frequency oscillations (HFOs) related to epilepsy ^73^, we discarded any candidate ripple event that occurred within 500 ms from inter-ictal epileptic discharges (IEDs). The detection of IEDs involved two complementary methods: an automated line-length algorithm based on ^74^ and a band-limited power analysis similar to ^75^. For the latter, the raw hippocampal iEEG signal was first filtered between 20-40Hz using a zero-lag linear-phase Hamming windowed FIR filter. Next, the filtered signal underwent rectification, squaring, smoothing, and z-scoring. Events that exceeded 8 standard deviations (SD) were identified as IEDs.

### Spectral analysis

Spectral decomposition of ripple events was done using a Morlet-wavelet time-frequency method, implemented in EEGLAB. We used a window of 2 cycle at the lowest frequency (4 Hz) and up to 20 cycles at the highest plotted frequency (220 Hz), with a step size of 6 ms. Ripple-triggered spectrograms were normalized by the geometric mean power in each frequency, computed over the entire epoch length (–750 to 750 ms) and across all epochs belonging to the same condition (i.e., rest (1), screening (1), rest (2), screening (2), memory test). The spectrograms of the videoclip responses were computed in EEGLAB using a Hanning windowed Fast Fourier Transform (FFT). The window size was 256 ms long, advanced with a step size of 1 ms, covering a frequency range from 4 to 128 Hz. These spectrograms were normalized by the geometric mean power in each frequency, computed over a pre-stimulus baseline period.

### Cross-correlation analysis

To evaluate the cross-correlation between the timing of ripple generation during the first and second screenings, we initially convolved the ripple train with a 4s Gaussian window (σ=0.25) and computed the cross-correlation using the xcorr.m function in Matlab. Normalization of the cross-correlation function was performed by dividing it by the mean cross-correlation values obtained from shuffled data. The shuffling was done by circularly shifting the ripple trains by a random amount 50 times and recalculating the cross-correlation. When assessing the cross-correlation using the video clip intervals, the null distribution was established by shuffling the clips 50 times.

### Statistical tests

All statistical analyses were performed in Matlab. Pairwise comparisons were carried out using two-sided Wilcoxon signed-rank and rank-sum tests, unless stated otherwise. We did not use statistical methods to predetermine sample sizes, however, sample sizes were similar to those generally employed in the field ^35,42,54,76^. Typically, the unit of analysis was individual patients. In cases where the units of analysis were individual electrodes, this was explicitly mentioned in the text. Resampling tests were carried out using either in-house MATLAB code or using established routines ^77^ implemented in the Mass Univariate ERP Toolbox ^78^. Multiple comparisons correction was carried out using either the false discovery rate adjustment (FDR) ^79^, or through cluster-based permutation tests ^49,77^.

### Mixed effects analysis

Mixed-effects analyses were carried out in MATLAB using the fitlme.m function. The data were fitted with a random intercept model that included the relevant fixed factors, as well as a random factor of ‘Electrode’ nested within ‘Patient’. The latter accounts for the fact that different participants contributed different numbers of electrodes to the analysis.

---

**Extended Data Figure 1.**
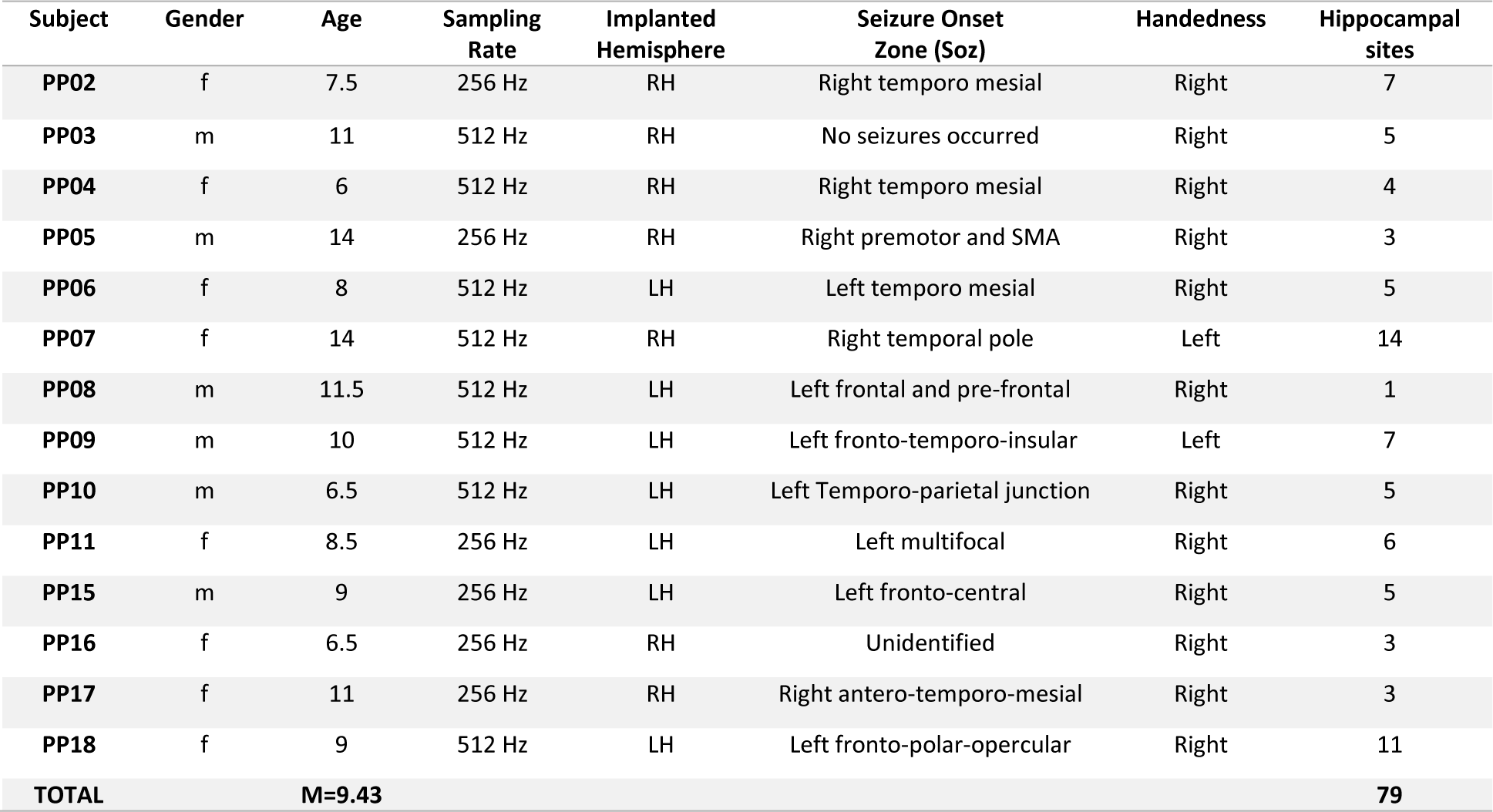
Demographic information, hippocampal electrode count and seizure onset zone for each patient.

**Extended Data Figure 2.**
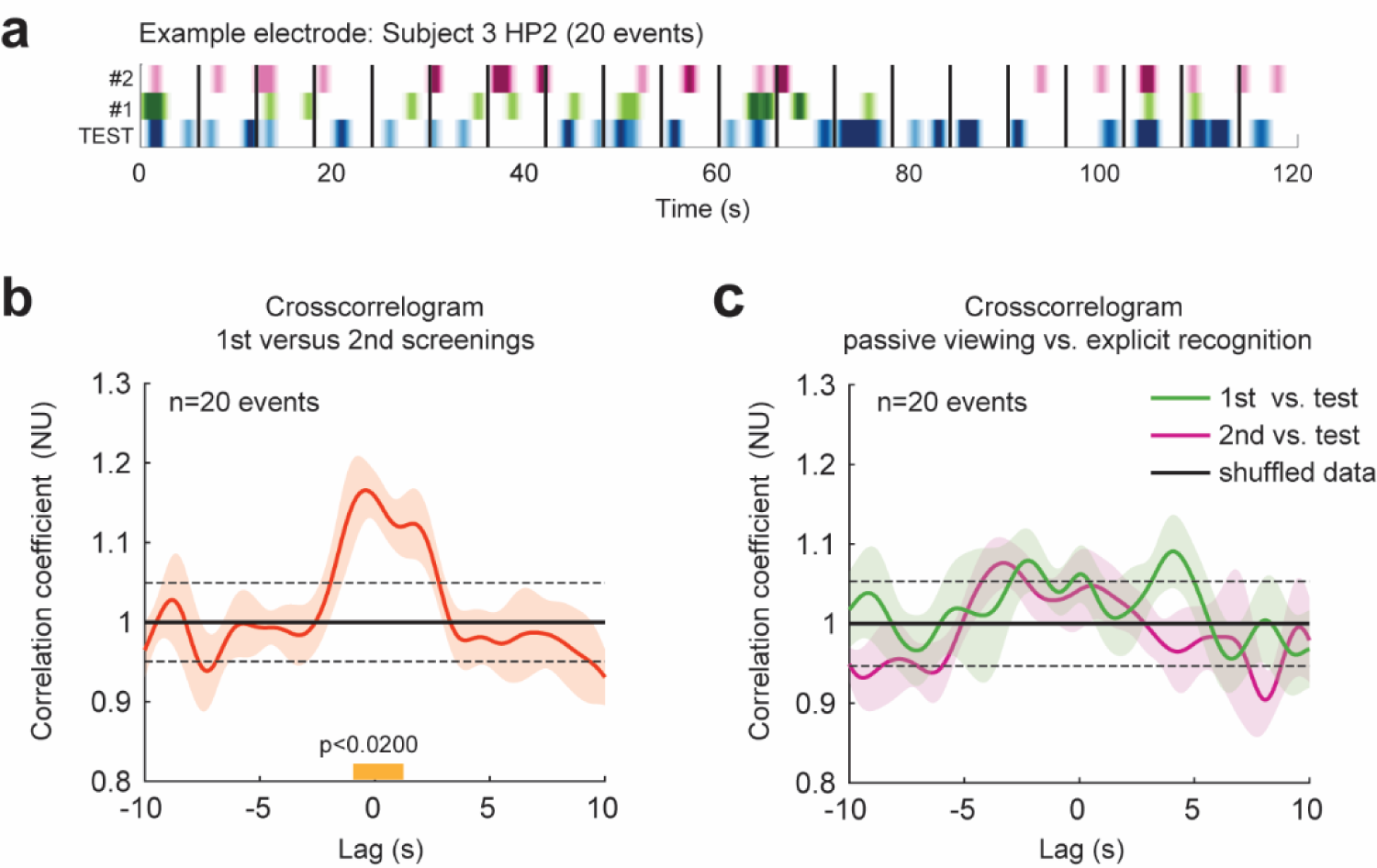
Ripples crosscorrelogram between the first and second screenings and the recognition test (a) An illustrative example of ripple train in one recording site during the 20 animated events that were extracted from the full-length screenings. The participants watched the same events under three viewing conditions: initial full-length screening (green), repeated full-length screening (pink) and memory test (blue). (b) Crosscorrelogram between the first and second screening of the cartoon, including only the intervals that corresponds to the videoclips. Similar to Figure 4f, there was a prominent peak around zero lag, highlighting the consistency in the temporal activation profile of ripples during the passive viewing of the cartoon. (c) Crosscorrelogram between the memory test and the first (green) and second (pink) screenings. The temporal activation profile of ripples during the recognition test did not correlate with the passive viewings of the cartoon. The black dashed line in (b) and (c) represents the mean±SD of cross-correlation magnitudes when the events were randomly shuffled 50 times.

